# The need for hemispheric separation in pairwise structural disconnection studies

**DOI:** 10.1101/2023.03.30.534883

**Authors:** Lisa Röhrig, Hannah Rosenzopf, Sofia Wöhrstein, Hans-Otto Karnath

## Abstract

The development of new approaches indirectly measuring the structural disconnectome has recently led to an increase in studies investigating pairwise structural disconnections following brain damage. Previous studies jointly analyzed patients with left hemispheric and patients with right hemispheric lesions when investigating a behavior of interest. An alternative approach would be to perform analyses separated by hemisphere, which has been applied in only a minority of studies to date. The present simulation study investigated whether joint or separate analyses (or both equally) are appropriate to reveal the ground truth disconnections. In fact, both approaches resulted in very different patterns of disconnection. In contrast to analyses separated by hemisphere, joint analyses introduced a bias to the disadvantage of intra-hemispheric disconnections. Intra-hemispheric disconnections were statistically underpowered in the joint analysis and thus surpassed the significance threshold with more difficulty compared to inter-hemispheric disconnections. This statistical imbalance was also shown by a greater number of significant inter-hemispheric than significant intra-hemispheric disconnections. This bias from joint analyses is based on mechanisms similar to those underlying the ‘partial injury problem’. We therefore conclude that pairwise structural disconnections in patients with unilateral left hemispheric and with unilateral right hemispheric lesions exhibiting a specific behavior (or disorder) of interest should be studied separately by hemisphere rather than in a joint analysis.

## Introduction

In the last years, the indirect estimation of structural disconnection has become an increasingly popular alternative to directly measured structural disconnectomes. The latter are available only from specific imaging sequences such as Diffusion Tensor Imaging (DTI), which are only sparsely available for most clinical cases. Software packages such as the Lesion Quantification Toolkit (LQT; Griffis et al., 2021), the Brain Connectivity and Behavior Toolkit (BCBtoolkit; Foulon et al., 2018), and the Network Modification (NeMo) tool (Kuceyeski et al., 2013) use lesion maps from neurological patients as seeds to estimate which fiber tracts intersect with the lesion, based on a tractography atlas of healthy individuals. In combination with a gray matter atlas, they further allow the creation of pairwise (also referred to as connection-wise, parcel-wise, region-wise, region-to-region, or ROI-to-ROI) disconnection measures between each gray matter region-pair.

Disconnection measures created this way have a decisive theoretical advantage over voxel-wise approaches in univariate statistics concerning the so-called ‘partial injury problem’ (Kinkingnéhun et al., 2007; Rorden et al., 2009; Sperber et al., 2019). The partial injury problem argues that (in voxel-wise designs) true lesion-deficit associations are deluded when lesions of different patients fall into different parts of the ground truth region (i.e., the entire area truly eliciting the behavioral deficit). In contrast, pairwise approaches are based on the summation of voxels according to predefined atlas parcels, so that a direct overlap of lesioned voxels is less essential for the successful discovery of an association. The creation of region-pairs even allows considering the mutual involvement of spatially distributed lesions due to a damaged connection between them, which further increases the statistical power in univariate analyses for pairwise disconnection measures compared to single voxels (see Sperber et al., 2022).

The mechanisms underlying the partial injury problem also apply to the pooled analysis of lesions in the left and right hemispheres. Even if structures relevant to a function of interest are represented in both hemispheres as it is the case for non-lateralized behaviors, lesion-symptom mapping that analyzes unilateral left and unilateral right lesions together causes voxels of either hemisphere to be statistically underpowered. Voxels of one hemisphere act as opponents for voxels of the other one. Therefore, individual analyses for each hemisphere have been established as the standard in voxel-wise lesion-symptom mapping (Frenkel-Toledo et al., 2019, 2021). However, so far it is unclear whether or not this problem applies also to pairwise disconnection approaches.

Recent literature addressing target behaviors in patients suffering from brain injury primarily analyzed structural disconnections in a joint analysis, i.e., a single analysis based on patients with unilateral left and with unilateral right hemispheric lesions (e.g., Errante et al., 2022; Kuceyeski et al., 2016; Moore et al., 2022; Pan et al., 2022; Smaczny et al., 2023; Urbanski et al., 2016), and only a minority in individual analyses separated by hemisphere (e.g., Rosenzopf et al., 2023). The question thus arises whether joint or separate analyses, or both equally, are valid to study pairwise structural disconnections in humans. Theoretically, the uneven distribution of inter-hemispheric disconnections (possible in the whole sample) and intra-hemispheric disconnections (possible only in patients with lesions in the respective hemisphere) might introduce a similar bias as known from lesions in opposing hemispheres (i.e., the partial injury problem). Joint analyses could underpower intra-hemispheric disconnections and decrease their chances to be uncovered (despite a true association to the deficit). Simulations offer an excellent approach to investigate these questions. We thus examined the effects of joint analyses versus analyses separated by hemisphere to answer the question whether it is more valid to analyze left and right hemispheric lesions jointly or separately in pairwise disconnection studies.

## Methods

All analyses were performed in MATLAB R2019a (The MathWorks, Inc., Natick, USA) via custom scripting. We conducted two simulation paradigms, for which we defined three pairwise ground truth connections: one inter-hemispheric connection and one intra-hemispheric connection in each hemisphere (see Fig. 1). Region-pairs were based on the Brainnetome Atlas (BNA; Fan et al., 2016). The inter-hemispheric connection extended between ventral area 9/46 of the middle frontal gyrus (MFG) of the left and right hemispheres; the intra-hemispheric connection extended between medial area 38 of the superior temporal gyrus (STG) and ventral agranular insular area of the insular gyrus of either hemisphere. To mimic a typical clinical cohort, we created samples that consist to one third of critical cases (i.e., patients with damage to a ground truth connection and the associated target behavior) and two thirds that represent ‘control’ cases.

**Figure 1.**
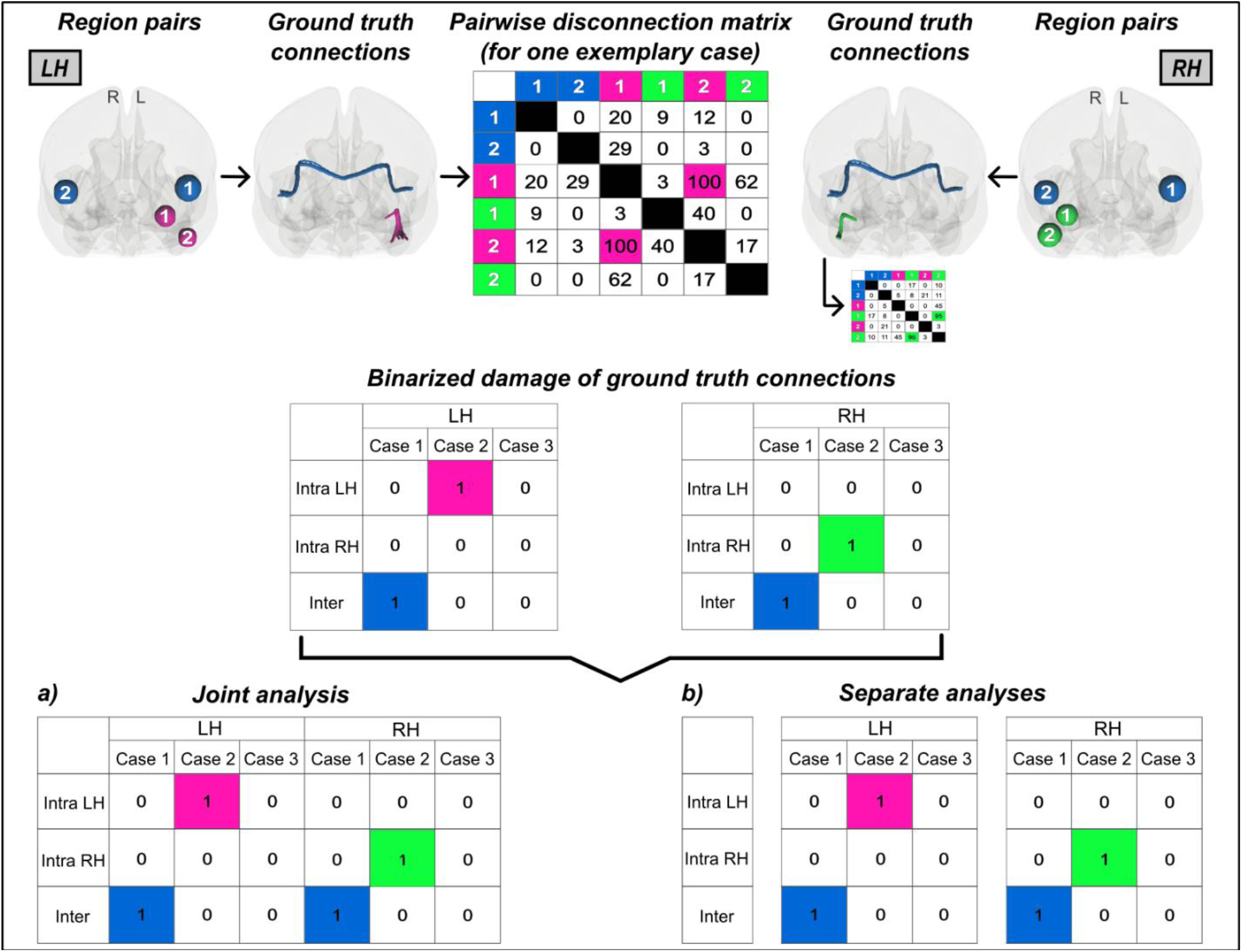
Ground truth connections and pairwise disconnection matrices. The three pairwise ground truth connections, whose damage was associated with the target behavior in the simulations, are visualized in the upper row for the left (‘LH’) and right hemisphere (‘RH’): one inter-hemispheric connection (blue) and two intra-hemispheric connections (LH: pink; RH: green). Shown are the corresponding region pairs schematically and the path of involved tracts of each ground truth connection based on the HCP-842 tractography atlas (Yeh et al., 2018) in anterior view using DSI studio (https://dsi-studio.labsolver.org/). The pairwise disconnection matrix of one exemplary case (with damage to the left intra-hemispheric connection) is depicted in a simplified manner; real matrices consisted of 246×246 cells. For simplicity, the continuous disconnection scores of the three ground truth connections are visualized as binary values for unilateral left and unilateral right brain damage for three exemplary cases each (middle section). The two analysis approaches of jointly or separately analyzing left and right hemispheric pairwise disconnections are illustrated at the bottom. This simplified illustration shows the difference in statistical power between inter- and intra-hemispheric disconnections for the two analysis approaches. Left and right hemispheric brain damage can only be unified by inter-hemispheric connections because these are the only ones they share. All intra-hemispheric connections remain hemisphere-specific entities that are subject to an issue similar to the ‘partial injury problem’.

### Simulation 1: Pairwise disconnection-based simulation

The first simulation served as the maximally controlled and purely methodological validation of both analysis approaches, in which we directly simulated the pairwise disconnection matrices for 50 cases per hemisphere. Here, we could design the disconnections as it is not possible in real-world scenarios and hence demonstrate the genuine statistical impact. For the left and right hemisphere separately, we created 246×246 symmetric matrices (simplified visualized in Fig. 1) in accordance with the gray matter parcellation atlas BNA that consists of 246 parcels based on structural and functional connections (Fan et al., 2016). Each matrix contained randomly generated continuous disconnection scores between 0 and 100 for randomly selected pairwise direct inter- and intra-hemispheric connections of one hemisphere. To avoid extremes, the number of disconnections was set to range between 50 and the value representing half of all possible direct connections of the respective hemisphere (i.e., 1181 in the left, 1111 in the right hemisphere). The remaining connections were considered unaffected and set to 0. For the control cases (*N* = 34), we set the disconnection of the three ground truths to 0. For the critical cases (*N*_*inter*_ = 8; *N*_*intra*_ = 8), we pseudo-randomly generated a disconnection score between 30 and 100 (considering a score greater than 30 as pathological) for either the inter- or the intra-hemispheric connection. Scores for the critical ground truth inter-hemispheric and intra-hemispheric connections were identical to ensure equal prerequisites. The simulated behavior was identical, with the exception that the range was set to 0 to 1, to the maximum disconnection score of the ground truth connections in the case-specific matrix. That means, damage to each of the three target connections was individually considered to lead to the target behavior, which was inspired by the equivalence ‘brain mode’ (see, e.g., Toba et al., 2020).

To investigate the association between structural disconnectivity and behavior (both continuous variables), we utilized mass-univariate pairwise analyses based on online available scripts (cf. Sperber et al., 2022). The simulated pairwise disconnection matrices were used; redundant cells and connections damaged in less than 5 cases were removed to reduce the number of tested connections by neglecting those with low variance. For each pairwise connection (end-to-end), we computed a General Linear Model (GLM). To correct for multiple testing, we ran 10,000 iterations with pseudo-randomly permuted behavior by implementing the maximum-statistics approach, for which in each permutation iteration the maximum t-statistic of all tested pairwise connections was saved. Among the distribution of those maximum t-statistics, the significance threshold was adjusted to get a family-wise error (FWE)-corrected *p* < 0.05.

### Simulation 2: Lesion-based simulation

In the second simulation, we pursued a more realistic approach by generating 3D lesion maps and indirectly estimating their corresponding pairwise disconnections. We generated 50 binary lesion maps per hemisphere as ellipsoids with random radiuses in standard MNI space. For each hemisphere, we created randomly distributed lesions, of which some were localized along the two targeted pairwise connections. After lesion map generation, maps were masked by ventricles, cerebellum, extracerebral space, and the opposite hemisphere at the midsagittal plane. The overlap of the simulated lesions is shown in Figure 2. Mean lesion volume was 13.6 ml (SD: 10.0 ml) in total, 13.4 ml (SD: 10.8 ml) for left-sided lesions, and 13.7 ml (SD: 9.3 ml) for right-sided lesions.

**Figure 2.**
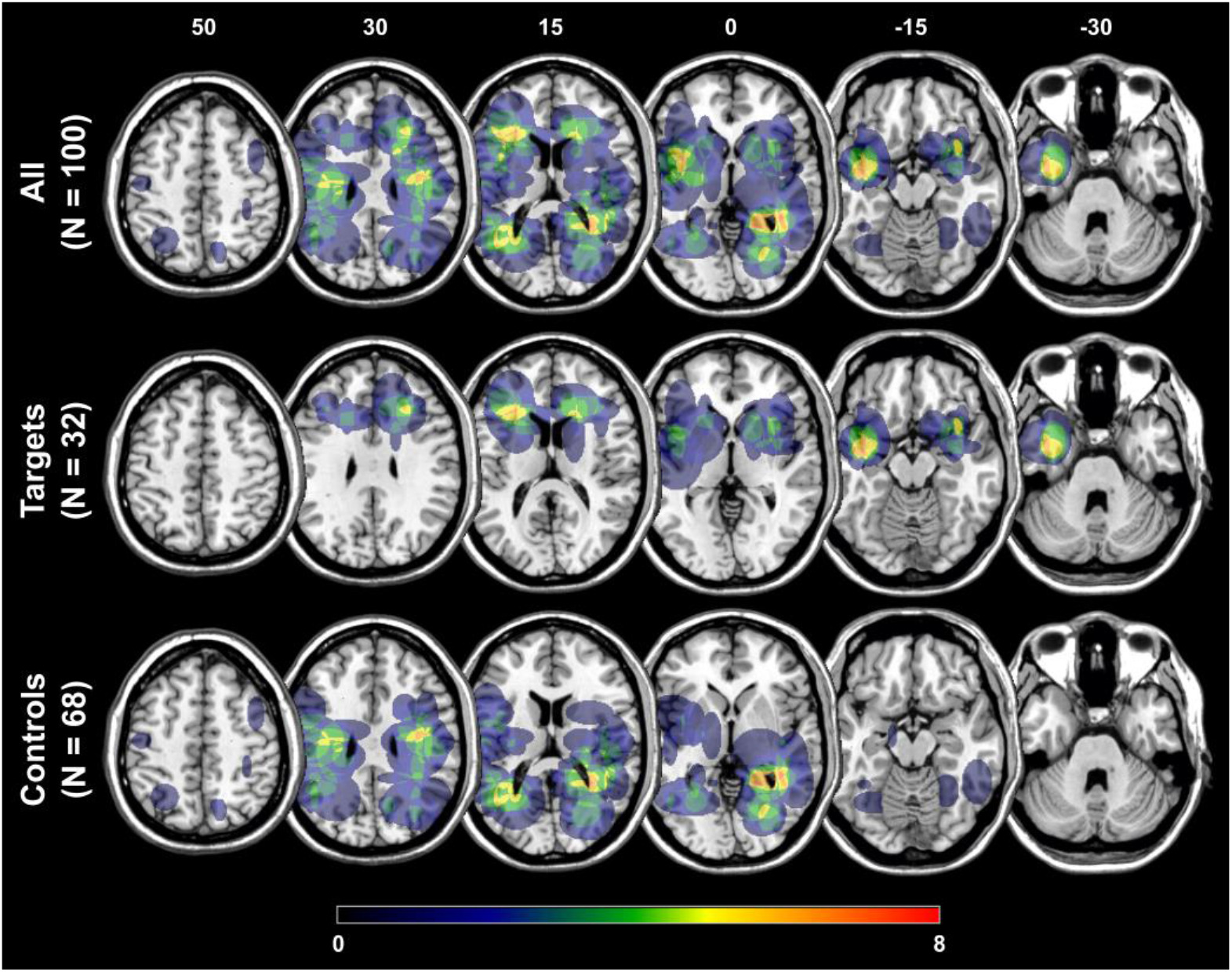
Overlap of the simulated lesions. Voxel-wise overlap of the simulated lesions (*N* = 50 per hemisphere) is shown for the whole sample (top row), critical target lesions (middle row), and ‘control’ lesions (bottom row) on the ch2-template in MNI space via MRIcron (https://www.nitrc.org/projects/mricron). Lesions were randomly generated for the left and right hemisphere separately. The color bar indicates the number of overlapping lesions for each voxel. Numbers above each column represent the z-coordinate (mm) in MNI space.

We then indirectly estimated the structural disconnectome: the generated binary lesion maps served as seeds to track all fibers passing through each lesion. The LQT with the default options was selected for this purpose. We used the default tractography atlas based on 842 healthy individuals by the Human Connectome Project (Yeh et al., 2018) and the BNA as gray matter parcellation atlas. For each simulated lesion, the obtained pairwise disconnection matrix contains the relative damage for each region-pair that has a direct anatomical connection according to the tractography atlas. In accordance with our first simulation, lesions with a pairwise disconnection of at least 30 percent for a target connection were defined as critical lesion leading to the behavior of interest; for each hemisphere, we generated 8 critical lesions with damage to the inter-hemispheric target connection and 8 critical lesions with damage to the intra-hemispheric target connection.

For the simulation of the target behavior, we again used the maximum pairwise disconnection score among the three ground truth connections (and set the range to 0 to 1). To simulate different natural scenarios of statistical power, we inserted a specified proportion of noise to the maximum disconnection score; we tested noise levels of 0, 10, 25, 45, 50, 55, and 75 percent of the total behavioral score. The noise value was pseudo-randomly defined and equal for inter- and intra-hemispheric critical subsamples to ensure that none of the target connections accidentally received overall more noise than the other that would lead to an unequal blurring of the connection-behavior association. The pseudo-random noise generation had the additional advantage that all simulation analyses (left and right hemispheric samples as well as different noise levels) were directly comparable. We then implemented mass-univariate analyses with permutation-based correction for multiple testing as explained above for the first simulation.

In addition, we looked at the effect of a greater sample size by extending the sample to 120 lesions per hemisphere (*N*_*control*_ = 80; *N*_*critical-inter*_ = 20; *N*_*critical-intra*_ = 20; for an overlay image, see Fig. S1 in the Supplement). The following procedure was similar to the one described above, except that we kept pairwise connections damaged in at least 20 cases and that we tested noise levels of 70 and 80 percent instead of 45 and 55 percent due to the increased statistical power.

## Results

In both the first and second simulations, the joint analysis and the individual analyses separated by hemisphere revealed very different pairwise disconnection patterns. In general, analyses separated by hemisphere showed a more realistic image with respect to the simulated connection-behavior association of the respective inter- or intra-hemispheric ground truth connections.

### Simulation 1: Pairwise disconnection-based simulation

When analyzing left and right hemispheric pairwise disconnections together, only the inter-hemispheric disconnection became significantly associated to the simulated behavior (*t* = 8.37; *p* < 0.0001; Fig. 3). This indicates that varying degrees of relevance were assigned to the association between inter- and intra-hemispheric disconnections to the target behavior, whereby the intra-hemispheric disconnections were not detected to be significantly associated to the target behavior (*t* = 4.83; *p* > 0.05). In contrast, when separating the analyses according to hemisphere, damage to both ground truth connections of each hemisphere, i.e., the inter-hemispheric as well as the intra-hemispheric connection, was significantly linked to the simulated behavior (*t* = 5.86; *p* < 0.05; Fig. 3). Interestingly, the obtained t-statistics for inter- and intra-hemispheric disconnections were identical, indicating that none of them were subject to a methodology-induced bias and in fact reflect reality, as the ground truth connections were indeed simulated as being equally relevant. Besides the ground truth connections, no further disconnections became significant.

**Figure 3.**
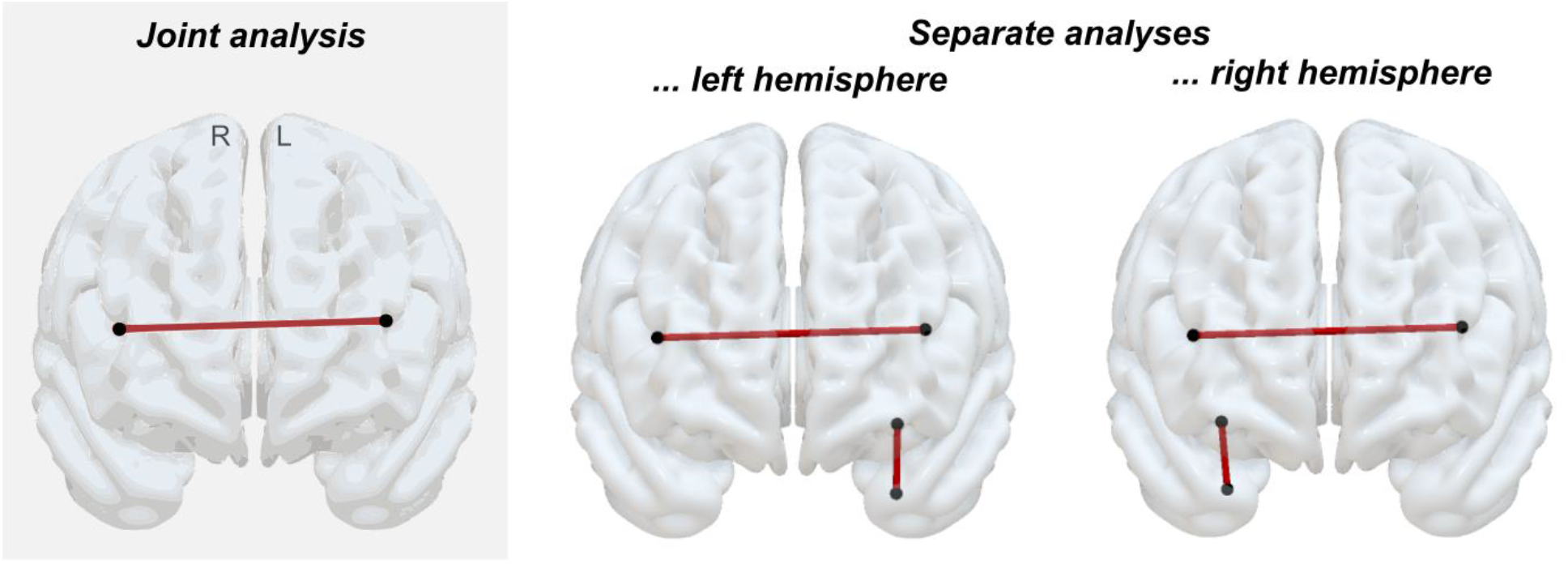
Significant pairwise disconnections obtained from the pairwise disconnection-based simulation. Disconnections significantly associated to the simulated behavior are shown in the anterior view (‘L’ – left hemisphere, ‘R’ – right hemisphere) using SurfIce (https://www.nitrc.org/projects/surfice/). Results are depicted for the joint analysis (left, gray shaded column) as well as for the individual analyses separated by hemisphere (*p* < 0.05, corrected for multiple comparisons). The inter-hemispheric ground truth disconnection turned out significant in both the joint and the separate analyses; the left and the right intra-hemispheric ground truth disconnections became significant in the analyses separated by hemisphere, but not in the joint analysis.

### Simulation 2: Lesion-based simulation

The obtained t-statistics for each ground truth connection and for the different tested noise levels are visualized in Fig. 4A. For the intra-hemispheric ground truth connections, the t-statistics are reduced in the joint analysis compared to the hemisphere-specific analysis, whereas for the inter-hemispheric ground truth connection, the t-statistic is enhanced in the joint analysis compared to the separate analyses. In more detail, the inter-hemispheric connection was remarkably more significant than both intra-hemispheric connections in the joint analysis, while in contrast, the intra-hemispheric connections were (slightly) more significant than the inter-hemispheric one in both individual analyses (see Fig. 4A). Hence, we observed a switched order of the most relevant disconnections between both analysis approaches. With increasing noise, the overall power of the ground truth connections decreased, which led to intra-hemispheric connections no longer surpassing the significance threshold in the joint analysis, as it is the case for 55 percent noise, for instance (see Fig. 4A). Looking at the numbers of significant inter- and intra-hemispheric disconnections (Fig. 4B and 4C), we found a large amount of significant inter-hemispheric disconnections compared to a (much) smaller amount of intra-hemispheric disconnections surpassing the significance threshold in the joint analysis across noise levels. The results obtained by the joint analysis approach suggest that mainly inter-hemispheric disconnections are associated with the behavior of interest. On the other hand, when investigating left and right hemispheric lesions and their corresponding disconnectomes in separate analyses, we found fewer significant inter-hemispheric disconnections than intra-hemispheric ones for left and right-sided lesions (Figure 4B and 4C).

**Figure 4.**
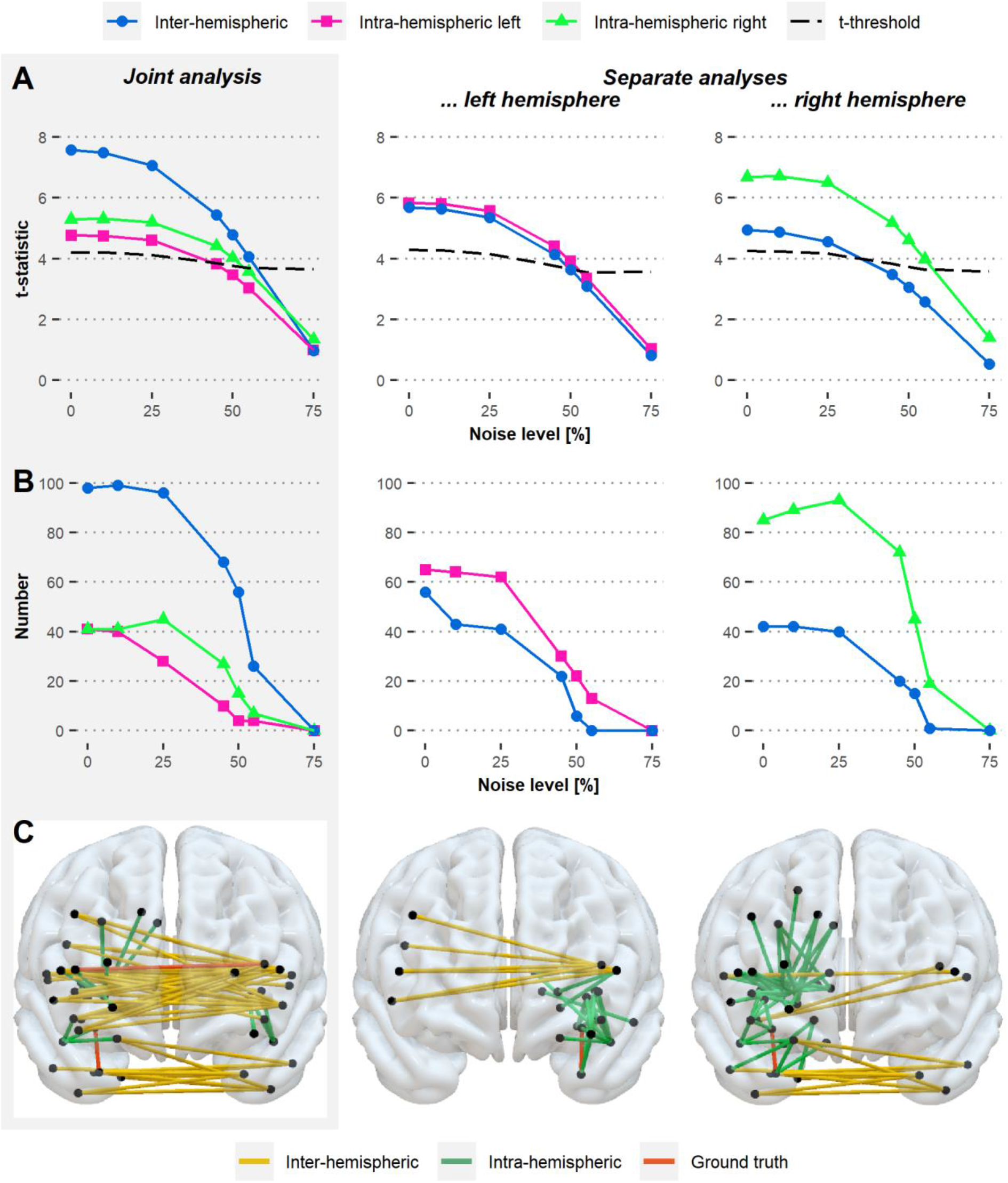
Effects of joint and of separate analysis approaches on pairwise disconnections induced by simulated lesions. Results were obtained from the lesion-based simulation for 50 lesions per hemisphere after permutation-based correction for multiple testing and are visualized for the joint analysis (left, gray shaded column) and analyses separated by hemisphere across different noise levels (i.e., 0, 10, 25, 45, 50, 55, and 75 percent). **(A)** For each pairwise ground truth connection, t-statistics are visualized. The dashed line represents the corrected significance threshold across the noise levels; t-statistics displayed above the line represent results significant at *p* < 0.05. (Note that the inter-hemispheric ground truth connection was truly less relevant compared to the intra-hemispheric one in both hemispheres owed to the simulated data, as visible for the zero-noise condition for the separate analyses.) **(B)** Numbers of inter- and intra-hemispheric disconnections surpassing the significance threshold (*p* < 0.05) are depicted. **(C)** The pattern of pairwise disconnections significant at *p* < 0.05 is shown in the anterior view exemplary for the 50 percent noise level.

When analyzing a much larger sample of 240 lesions in total, we found similar effects as the ones described above for the two investigated analysis approaches (see Fig. S2 in the Supplement). Greater statistical power due to a larger sample size does therefore not prevent from the statistical bias induced by the joint analysis. However, the statistical power did benefit from an increased sample size so that certain ground truth connections could be detected in the separate analyses for noise levels, for which those connections could not be detected by the separate analysis approach using the smaller sample (e.g., 50 percent noise level).

## Discussion

In the present simulation study, we tested whether a single analysis of both unilateral left and unilateral right hemispheric lesions together or analyses separated by hemisphere would result in comparable pairwise, region-to-region structural disconnection results, or whether one or the other analysis approach should be preferred. We associated behavior with pairwise structural disconnections, which were either (i) directly simulated or (ii) estimated indirectly from simulated unilateral left and unilateral right hemispheric lesions. We observed that joint and separate analyses resulted in very different patterns of disconnection. Analyses separated by hemisphere showed a more realistic image with respect to the simulated connection-behavior association of the respective inter- or intra-hemispheric ground truth connections.

In the joint analysis of left and right hemispheric lesions, intra-hemispheric disconnections were statistically underpowered and thus surpassed the significance threshold with more difficulty compared to inter-hemispheric disconnections. In contrast, analyses separated by hemisphere did not neglect one type of disconnection. Our first, pairwise disconnection-based simulation, which represents a scenario of perfect circumstances, clearly preferred the approach of analyzing both hemispheres separately, because only this approach could detect all ground truth connections. This finding indicates that the procedure of separate analyses is methodologically seen the undisputedly best approach. Our second simulation, which represents the noisier but more natural scenario, showed similar effects. Different from our first simulation, with increasing noise, the separate analyses as well as the joint analysis could not detect all ground truth connections. However, this issue is more of a general problem resulting from lower effect sizes due to insufficient sample sizes. Our simulation revealed that – at the same noise level – an increased sample size led to the detection of all underlying ground truth connections in the separate analysis approach, whereas the statistical bias was still preserved in the joint analysis. Our data thus suggest that – depending on different effect sizes, sample sizes, and background noise of the respective data set − the sole application of analyses separated by hemisphere can already be sufficient, while in other cases, separate analyses might not catch all ground truth disconnections because of the statistical threshold used and the insufficiently large effect size of the missed disconnection. However, increasing the sample size might eliminate this issue as shown by the present simulation. Also, reporting the uncorrected results might be an option to deal with small sample sizes.

In a recent article of our group, we conducted both a joint analysis of unilateral left and right hemispheric patients as well as individual analyses separated by hemisphere on real data of stroke patients suffering from ‘pusher syndrome’ (Rosenzopf et al., 2023). We observed that the results obtained from the respective analyses indeed were in accordance with the here described pattern. Indeed, we obtained a smaller amount of significant results when analyzing both hemispheres together; all additional disconnections that appeared in the separate analyses were ipsilesional intra-hemispheric disconnections (Rosenzopf et al., 2023).

Besides the statistical bias that results to the disadvantage of intra-hemispheric disconnections, there are further aspects arguing against the application of joint analyses. First, even if all ground truth connections would survive the significance threshold due to sufficiently large effect sizes, this analysis approach still impacts the level of significance of disconnections. Intra-hemispheric disconnections would still be misinterpreted in a joint analysis as less relevant than inter-hemispheric ones. In fact, they might be ignored since most researchers tend to discuss only the most significant findings, especially when there are dozens of significant items. Second, another argument against joint analyses is that findings are less interpretable with respect to intra-hemispheric disconnections, since a significant intra-hemispheric disconnection does not allow any meaningful conclusion about the whole sample (i.e., the behavior/deficit of interest) because such a disconnection could only be caused by part of the sample.

Sperber et al. (2022) concluded that the partial injury problem is less of an issue in disconnection studies since pairwise measures can sum up damage that affects the same fiber tract or pairwise connection regardless of distance between them. While this is true for the ground truth correlate of a single hemisphere, our present results demonstrated that this conclusion is not valid for ground truth correlates located within both hemispheres. We have shown that mechanisms similar to those that cause power limitations in voxel-wise mapping of lesion-symptom-associations also operate in pairwise disconnection studies if the sample consists of lesions in both hemispheres. Joint analyses decrease the probability of intra-hemispheric disconnections truly associated with a behavior of interest to be uncovered. Inter-hemispheric connections can be damaged in both left and right-brain damaged patients, while, on the other hand, (direct) intra-hemispheric connections can only be disturbed within the lesioned hemisphere. In short, the necessary statistical power might not be achieved for intra-hemispheric disconnections when analyzing left- and right-sided lesions together, while inter-hemispheric disconnections profit from the increased sample size. An exclusively performed joint analysis can therefore solely make specific inferences on the potential involvement of inter-hemispheric disconnections while systematically underestimating possibly existing intra-hemispheric disconnections. Analyses separated by hemisphere, on the other hand, seem to reflect the true associations more validly. The statistical power in this approach might be overall lower than in the joint analysis due to the smaller sample size and potentially smaller effect size of a ground truth connection. However, neither the power of inter-hemispheric nor intra-hemispheric disconnections is systematically biased in analyses separated by hemisphere. The latter approach therefore does not favor the results for one connection type. The obtained t-statistics of the ground truth connections of either hemisphere were closer together in the individual analyses compared to the joint analysis, reflecting the simulated data in a more realistic manner.

Since the systematic bias induced by the joint analysis is a consequence of the sampling rather than the applied statistical approach, it would also persist in multivariate approaches and Bayesian frameworks, which are known to alleviate some issues associated with univariate approaches and frequentist statistics, respectively. In the joint analysis, inter-hemispheric disconnections would receive larger Bayes factors than intra-hemispheric disconnections, reflecting the same mechanisms described in the present simulation using t-statistics. Moreover, the present conclusions retrieved from pairwise structural disconnection-deficit-associations by unilateral brain lesions, such as strokes or tumors, are very likely to also apply to tract-wise disconnection measures, which define the damage not based on disconnected region-pairs but on the disconnected white matter fiber tracts. Tracts, like region pairs, can be characterized as inter-hemispheric (when the tract passes through both hemispheres) or intra-hemispheric (when the tract remains within one hemisphere) and are therefore likely susceptible to the same statistical bias we found here for pairwise measures. However, future studies should verify this assumption.

## Conclusion

The results derived from the present simulation study on pairwise structural disconnection analyses indicate that jointly analyzing unilateral left and unilateral right hemispheric lesions leads to a methodological bias, whereas the approach of individual analyses separated by hemisphere represents the accurate way to deal with such data sets. Joint analyses systematically underestimate possibly existing intra-hemispheric disconnections, which might therefore be considered less relevant or even completely missed. In contrast, the approach of hemispheric separation avoids such biased results. Hence, we propose that analyses separated by hemisphere should be considered as standard for pairwise (and tract-wise) structural disconnection studies.

## Acknowledgments

This work was supported by the Deutsche Forschungsgemeinschaft (KA 1258/23-1).

## Data availability statement

Simulated data and analysis results are openly available at Mendeley Data (DOI: 10.17632/rwgyvnsbm7.1).

## References

Errante, A., Rossi Sebastiano, A., Ziccarelli, S., Bruno, V., Rozzi, S., Pia, L., Fogassi, L., & Garbarini, F. (2022). Structural connectivity associated with the sense of body ownership: A diffusion tensor imaging and disconnection study in patients with bodily awareness disorder. Brain Communications, 4(1), fcac032. https://doi.org/10.1093/braincomms/fcac032

Fan, L., Li, H., Zhuo, J., Zhang, Y., Wang, J., Chen, L., Yang, Z., Chu, C., Xie, S., Laird, A. R., Fox, P. T., Eickhoff, S. B., Yu, C., & Jiang, T. (2016). The Human Brainnetome Atlas: A New Brain Atlas Based on Connectional Architecture. Cerebral Cortex, 26(8), 3508–3526. https://doi.org/10.1093/cercor/bhw157

Foulon, C., Cerliani, L., Kinkingnéhun, S., Levy, R., Rosso, C., Urbanski, M., Volle, E., & Thiebaut de Schotten, M. (2018). Advanced lesion symptom mapping analyses and implementation as BCBtoolkit. GigaScience, 7(3), giy004. https://doi.org/10.1093/gigascience/giy004

Frenkel-Toledo, S., Fridberg, G., Ofir, S., Bartur, G., Lowenthal-Raz, J., Granot, O., Handelzalts, S., & Soroker, N. (2019). Lesion location impact on functional recovery of the hemiparetic upper limb. PLOS ONE, 14(7), e0219738. https://doi.org/10.1371/journal.pone.0219738

Frenkel-Toledo, S., Ofir-Geva, S., Mansano, L., Granot, O., & Soroker, N. (2021). Stroke Lesion Impact on Lower Limb Function. Frontiers in Human Neuroscience, 15. https://www.frontiersin.org/articles/10.3389/fnhum.2021.592975

Griffis, J. C., Metcalf, N. V., Corbetta, M., & Shulman, G. L. (2021). Lesion Quantification Toolkit: A MATLAB software tool for estimating grey matter damage and white matter disconnections in patients with focal brain lesions. NeuroImage: Clinical, 30, 102639. https://doi.org/10.1016/j.nicl.2021.102639

Kinkingnéhun, S., Volle, E., Pélégrini-Issac, M., Golmard, J.-L., Lehéricy, S., du Boisguéheneuc, F., Zhang-Nunes, S., Sosson, D., Duffau, H., Samson, Y., Levy, R., & Dubois, B. (2007). A novel approach to clinical–radiological correlations: Anatomo-Clinical Overlapping Maps (AnaCOM): Method and validation. NeuroImage, 37(4), 1237–1249. https://doi.org/10.1016/j.neuroimage.2007.06.027

Kuceyeski, A., Maruta, J., Relkin, N., & Raj, A. (2013). The Network Modification (NeMo) Tool: Elucidating the Effect of White Matter Integrity Changes on Cortical and Subcortical Structural Connectivity. Brain Connectivity, 3(5), 451–463. https://doi.org/10.1089/brain.2013.0147

Kuceyeski, A., Navi, B. B., Kamel, H., Raj, A., Relkin, N., Toglia, J., Iadecola, C., & O’Dell, M. (2016). Structural connectome disruption at baseline predicts 6-months post-stroke outcome. Human Brain Mapping, 37(7), 2587–2601. https://doi.org/10.1002/hbm.23198

Moore, M. J., Hearne, L., Demeyere, N., & Mattingley, J. (2022). Comprehensive voxel-wise, tract-based and network lesion mapping reveals unique architectures of right and left visuospatial neglect. PsyArXiv. https://doi.org/10.31234/osf.io/jxrv4

Pan, C., Li, G., Jing, P., Chen, G., Sun, W., Miao, J., Wang, Y., Lan, Y., Qiu, X., Zhao, X., Mei, J., Huang, S., Lian, L., Wang, H., Zhu, Z., & Zhu, S. (2022). Structural disconnection-based prediction of poststroke depression. Translational Psychiatry, 12(1), Article 1. https://doi.org/10.1038/s41398-022-02223-2

Rorden, C., Fridriksson, J., & Karnath, H.-O. (2009). An evaluation of traditional and novel tools for lesion behavior mapping. NeuroImage, 44(4), 1355–1362. https://doi.org/10.1016/j.neuroimage.2008.09.031

Rosenzopf, H., Klingbeil, J., Wawrzyniak, M., Röhrig, L., Sperber, C., Saur, D., & Karnath, H.-O. (2023). Thalamocortical disconnection involved in pusher syndrome. Brain, awad096. https://doi.org/10.1093/brain/awad096

Smaczny, S., Sperber, C., Jung, S., Moeller, K., Karnath, H. O., & Klein, E. (2023). Disconnection in a left-hemispheric temporo-parietal network impairs multiplication fact retrieval. NeuroImage, 268, 119840. https://doi.org/10.1016/j.neuroimage.2022.119840

Sperber, C., Griffis, J., & Kasties, V. (2022). Indirect structural disconnection-symptom mapping. Brain Structure and Function, 227(9), 3129–3144. https://doi.org/10.1007/s00429-022-02559-x

Sperber, C., Wiesen, D., & Karnath, H.-O. (2019). An empirical evaluation of multivariate lesion behaviour mapping using support vector regression. Human Brain Mapping, 40(5), 1381–1390. https://doi.org/10.1002/hbm.24476

Toba, M. N., Godefroy, O., Rushmore, R. J., Zavaglia, M., Maatoug, R., Hilgetag, C. C., & Valero-Cabré, A. (2020). Revisiting ‘brain modes’ in a new computational era: Approaches for the characterization of brain-behavioural associations. Brain, 143(4), 1088–1098. https://doi.org/10.1093/brain/awz343

Urbanski, M., Bréchemier, M.-L., Garcin, B., Bendetowicz, D., Thiebaut de Schotten, M., Foulon, C., Rosso, C., Clarençon, F., Dupont, S., Pradat-Diehl, P., Labeyrie, M.-A., Levy, R., & Volle, E. (2016). Reasoning by analogy requires the left frontal pole: Lesion-deficit mapping and clinical implications. Brain, 139(6), 1783–1799. https://doi.org/10.1093/brain/aww072

Yeh, F.-C., Panesar, S., Fernandes, D., Meola, A., Yoshino, M., Fernandez-Miranda, J. C., Vettel, J. M., & Verstynen, T. (2018). Population-averaged atlas of the macroscale human structural connectome and its network topology. NeuroImage, 178, 57–68. https://doi.org/10.1016/j.neuroimage.2018.05.027

