## Supplementary Material for "The need for hemispheric separation in pairwise structural disconnection studies"

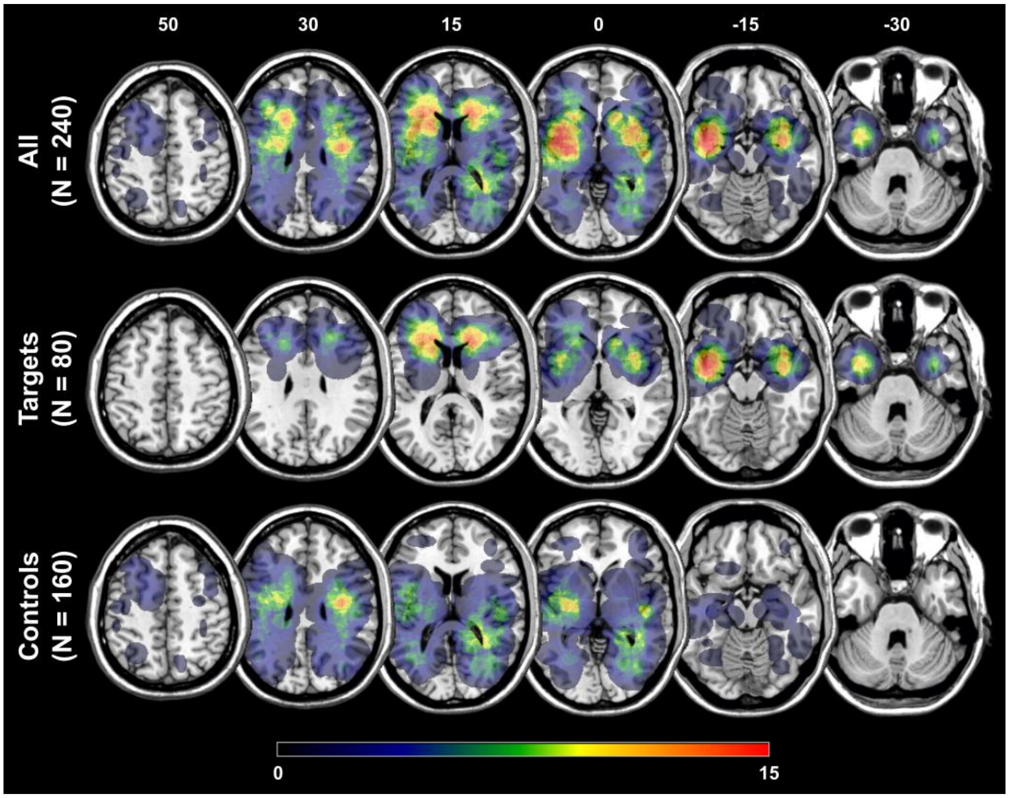


**Figure S1. Overlap of the 240 simulated lesions.** Voxel-wise overlap of the simulated lesions (*N* = 120 per hemisphere) is shown for the whole sample (top row), critical target lesions (middle row), and 'control' lesions (bottom row) on the ch2-template in MNI space via MRIcron (https://www.nitrc.org/projects/mricron). Lesions were randomly generated for the left and right hemisphere separately. The color bar indicates the number of overlapping lesions for each voxel. Numbers above each column represent the z-coordinate (mm) in MNI space. All lesions of the 50-sample (see main manuscript) are also included in the 120-sample.

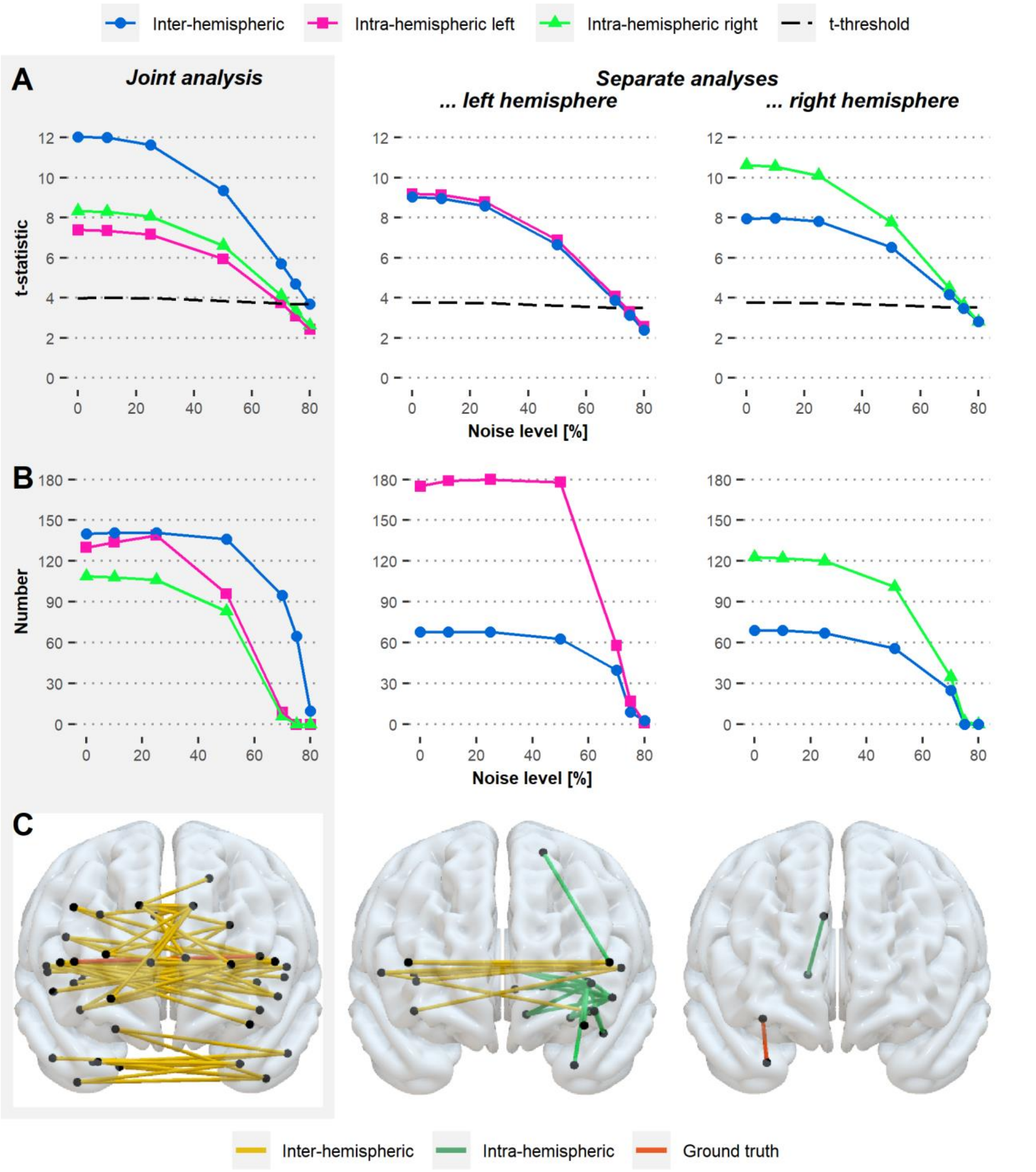


**Figure S2. Effects of joint and separate analysis approaches on pairwise disconnections induced by 240 simulated lesions.** Results were obtained from the lesion-based simulation for 120 lesions per hemisphere after permutation-based correction for multiple testing and are visualized for the joint analysis (left, gray shaded column) and analyses separated by hemisphere across different noise levels (i.e., 0, 10, 25, 50, 70, 75, and 80 percent). **(A)** For each pairwise ground truth connection, t-statistics are visualized. The dashed line represents the corrected significance threshold across the noise levels; t-statistics displayed above the line represent results significant at $p < 0.05$. (Note that the inter-hemispheric ground truth connection was truly less relevant compared to the intra-hemispheric one in both hemispheres owed to the simulated data, as visible for the zero-noise condition for the separate analyses.) **(B)** Numbers of inter- and intra-hemispheric disconnections surpassing the significance threshold ($p < 0.05$) are depicted. **(C)** The pattern of pairwise disconnections significant at $p < 0.05$ is shown in the anterior view exemplary for the 75 percent noise level.
